# Direction-selective motion discrimination by traveling waves in visual cortex

**DOI:** 10.1101/2020.03.03.974089

**Authors:** Stewart Heitmann, G. Bard Ermentrout

**Affiliations:** Victor Chang Cardiac Research Institute, Sydney, NSW, Australia; Department of Mathematics, University of Pittsburgh, PA, USA

## Abstract

The majority of neurons in primary visual cortex respond selectively to bars of light that have a specific orientation and move in a specific direction. The spatial and temporal responses of such neurons are non-separable. How neurons accomplish that computational feat without resort to explicit time delays is unknown. We propose a novel neural mechanism whereby visual cortex computes non-separable responses by generating endogenous traveling waves of neural activity that resonate with the space-time signature of the visual stimulus. The spatiotemporal characteristics of the response are defined by the local topology of excitatory and inhibitory lateral connections in the cortex. We simulated the interaction between endogenous traveling waves and the visual stimulus using spatially distributed populations of excitatory and inhibitory neurons with Wilson-Cowan dynamics and inhibitory-surround coupling. Our model reliably detected visual gratings that moved with a given speed and direction provided that we incorporated neural competition to suppress false motion signals in the opposite direction. The findings suggest that endogenous traveling waves in visual cortex can impart direction-selectivity on neural responses without resort to explicit time delays. They also suggest a functional role for motion opponency in eliminating false motion signals.

**Author summary:** It is well established that the so-called ‘simple cells’ of the primary visual cortex respond preferentially to oriented bars of light that move across the visual field with a particular speed and direction. The spatiotemporal responses of such neurons are said to be non-separable because they cannot be constructed from independent spatial and temporal neural mechanisms. Contemporary theories of how neurons compute non-separable responses typically rely on finely tuned transmission delays between signals from disparate regions of the visual field. However the existence of such delays is controversial. We propose an alternative neural mechanism for computing non-separable responses that does not require transmission delays. It instead relies on the predisposition of the cortical tissue to spontaneously generate spatiotemporal waves of neural activity that travel with a particular speed and direction. We propose that the endogenous wave activity resonates with the visual stimulus to elicit direction-selective neural responses to visual motion. We demonstrate the principle in computer models and show that competition between opposing neurons robustly enhances their ability to discriminate between visual gratings that move in opposite directions.

## Introduction

Hubel and Wiesel [1–3] laid the theoretical groundwork for the visual system with their discovery that many neurons in the primary visual cortex respond selectively to bars of light that move with a specific direction and speed. Such neurons are said to have non-separable space-time receptive fields (Fig 1A) because their responses to changing patterns of light and dark in the visual field cannot be explained in terms of independent spatial and temporal neural processes [4, 5]. The neural mechanism for computing non-separable responses is still an open question. Most theoretical accounts follow the approach of Reichardt [6] where light receptors exploit transmission delays to act as coincidence detectors of temporally delayed signals from disparate regions of the visual field (Figure 1B). The temporal delay essentially transforms the spatiotemporal computation into a spatial computation that can be feasibly accommodated by the dendritic arbors of a neuron. The concept was originally applied to direction-selective cells in the retina [7] and has since been extended to the visual cortex where transmission delays have been posited in the feed-forward projections [8–13] and in the lateral connections [14–16].

**Fig 1.**
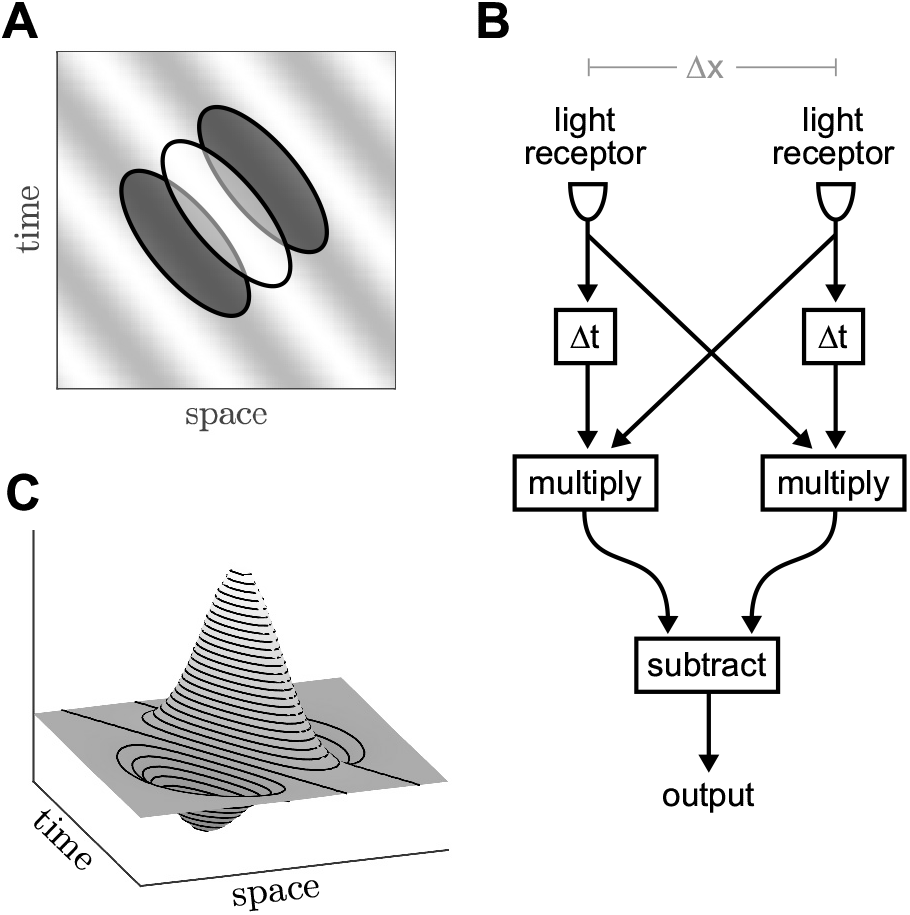
Spatiotemporal receptive fields. A: Schematic of a non-separable receptive field for a direction-selective neuron. The background grating represents a moving stimulus. The ellipses indicate the regions of light and dark that trigger a response in the neuron. B: Simplified schematic of the Reichardt [6] motion detector where Δ*x* is the spatial separation between light receptors and Δ*t* is a transmission delay. C: Gabor spatiotemporal filter constructed from a difference of Gaussians. Phase shifts are obtained by rotating the Gabor function in the space-time coordinate frame.

The motion-energy model [17] is a notable exception in that it uses the phase difference between time-varying signals in place of transmission delays. It accurately represents the responses of direction-selective neurons by applying a Gabor function (Fig 1C) to the features in the visual field [18, 19]. The Gabor function is constructed from a difference of Gaussians [20] that reasonably approximate the synaptic footprints of excitatory and inhibitory neurons. Yet the phase difference is imposed by rotating the Gabor function in the space-time coordinate frame without biophysical justification. Furthermore, the neural computation is expressed in terms of the visual coordinate frame rather than inputs at the neuronal level. Hence the motion-energy model is a descriptive model rather than an explanatory one [21].

Here we propose a neural mechanism for computing non-separable receptive fields without resort to explicit transmission delays. Our proposal relies on the retinotopic mapping of the visual system onto cortical coordinates and the propensity of cortical tissue to generate propagating waves of neural activity endogenously. We argue that those endogenous waves resonate with the spatiotemporal signature of stimulus to amplify the neural response for visual motion in a given speed and direction. It is those amplified responses that correspond to visual perception. The endogenous waves thus influence the responses of individual neurons and imbue them with their directional-selectivity. Furthermore, their preferred spatial and temporal frequencies are dictated by the geometry of the lateral inhibitory-surround coupling between excitatory and inhibitory neurons in the cortical tissue. This type of coupling features in the standard model of orientation selectivity that was originally proposed by Hubel and Wiesel [2] to explain the responses of ‘simple’ cells in primary visual cortex. Inhibitory-surround coupling also has conceptual links to the difference of Gaussians in the motion-energy model [17] and is crucial for the formation of standing wave patterns in neural field models [22–26].

Propagating waves have been observed in many regions of the brain [27–29] including the visual cortex. Stimulus-evoked and endogenously generated traveling waves have been observed in the visual cortex of monkey [30–34], cat [30, 35–37], rabbit [38], rat [39] and turtle [40]. The endogenous waves follow reproducible patterns that are related to the underlying anatomical connectivity [33, 41]. In cat visual cortex, those patterns are closely aligned with the functional orientation maps [37]. In human visual cortex, endogenous waves are thought to be the basis of geometric visual hallucinations for similar reasons [42, 43]. Stimulus-induced waves likewise follow reproducible patterns. Those waves travel beyond the footprint of the feed-forward projections [31] and are sensitive to the properties of the stimulus [30, 32, 44, 45]. More recently, Townsend and colleagues [34] found that the direction of waves elicited in primate visual cortex by drifting visual gratings and dot-fields are sensitive to the direction of the stimulus on a trial by trial basis. That particular study established a functional link between visual motion processing and traveling waves. It also demonstrated that sensory information can be encoded in cortical traveling waves at appropriate time scales. However the authors did not propose a neural mechanism to explain how that might be achieved.

In the present study, we used neural field models of the visual cortex to investigate how endogenous traveling waves interact with visual stimuli. Neural fields represent the large-scale activity of neural tissue as a spatial continuum where thousands of co-located neurons are lumped into localized populations called neural masses [46]. They are typically formulated in terms of the average membrane voltage or the average firing rate activity of the neurons [47]. Crucially for this study, neural fields produce spontaneous spatiotemporal patterns — called Turing patterns — under appropriate coupling conditions [22–24, 26, 48, 49]. In particular, Wilson and Cowan [48, 49] demonstrated standing wave patterns in a neural field with short-range excitatory connections and long-range inhibitory connections. Amari [22] later proved that result analytically for neural masses with a step-function firing-rate response and ‘Mexican hat’ coupling with distance. Such coupling topologies can be constructed from excitatory and inhibitory connection densities with Gaussian spatial profiles [50] and have direct analogy with the inhibitory-surround receptive fields described by Hubel and Wiesel [2].

We therefore modeled the visual cortex as a spatial continuum of excitatory and inhibitory neural populations with Gaussian coupling profiles where the spread of the inhibitory coupling exceeded that of the excitatory coupling by a factor of 3:1. We restricted our model to one spatial dimension for simplicity. The model produced endogenous standing wave patterns consistent with neural fields having ‘Mexican hat’ coupling [51, 52]. We then applied a small spatial shift to the profile of the excitatory connections to cause those standing waves to propagate with a given direction and speed. That asymmetric coupling was key to imbuing the medium with non-separable spatiotemporal response properties. We then explored how those endogenous waves resonated with drifting grating stimuli to elicit robust direction-selective responses to visual motion.

## Results

We defined the generalized equations for the Wilson-Cowan model (Fig 2A) as,

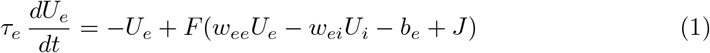

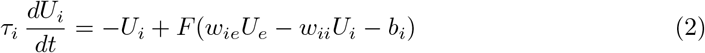

where *U_e_*(*t*) and *U_i_*(*t*) are the normalized firing rates of the excitatory and inhibitory neural populations. Both populations are reciprocally coupled where *w_ei_* denotes the weight of the connection from the inhibitory population to the excitatory population. The sigmoidal firing-rate function (Fig 2B) defines the response of each neural population to its input. Parameters *b_e_* and *b_i_* are the population firing thresholds. *J* (*t*) is an external stimulus which is applied to the excitatory population only.

**Fig 2.**
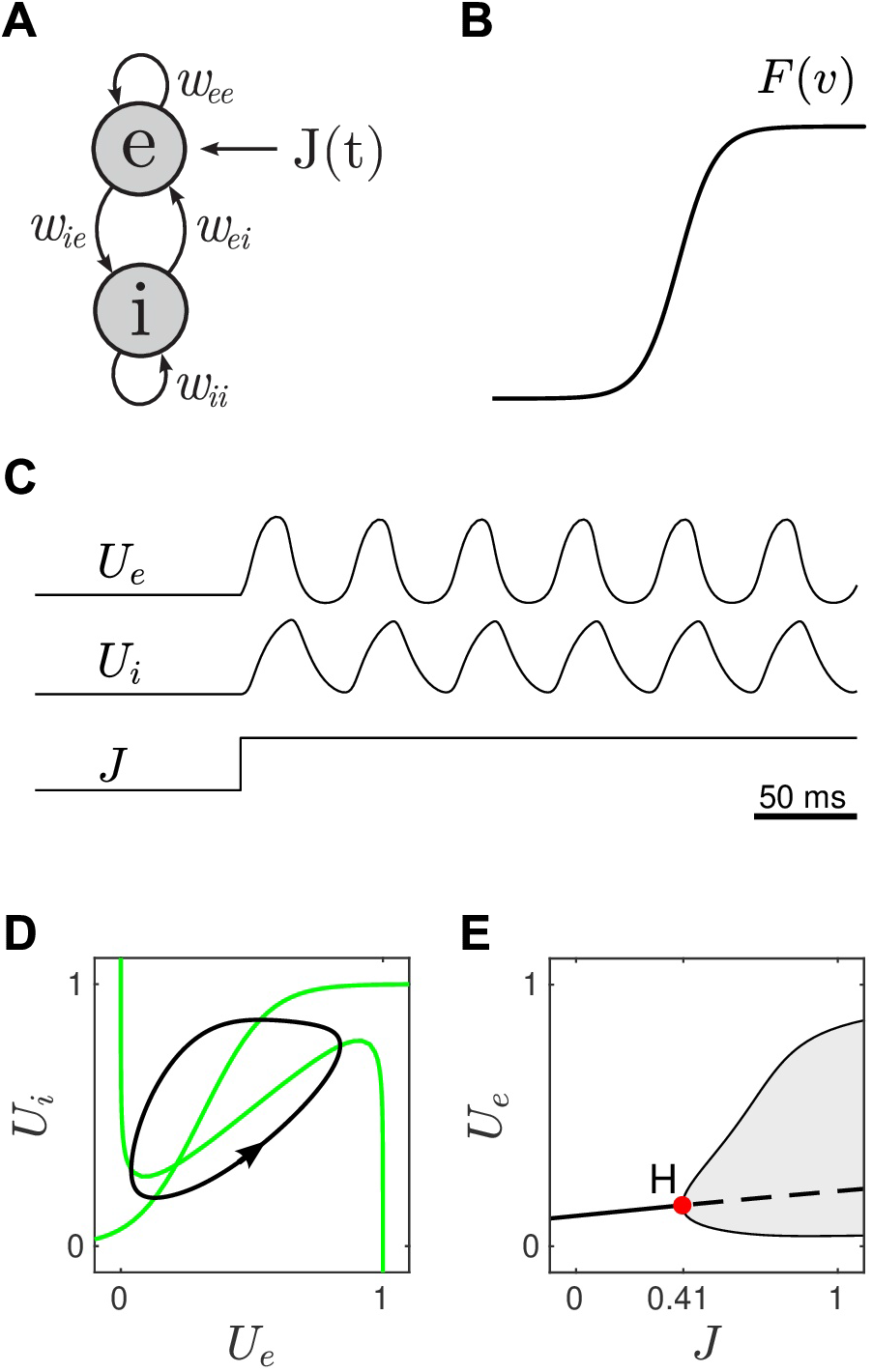
The Wilson-Cowan model of reciprocally-coupled excitatory and inhibitory neural populations. A: Schematic of the coupling where the weight for the connection to *e* from *i* is denoted by *w_ei_*. *J* (*t*) is an external stimulus. B: The sigmoidal firing rate function. C: Time course of the mean firing rates *U* (*t*) for both the excitatory and inhibitory populations in response to a unit-step stimulus. D: The limit cycle (black) in the phase plane. Nullclines are shown in green. E: Bifurcation diagram showing the emergence of the limit cycle (shaded region) via a supercritical Hopf bifurcation. Solid lines indicate stable fixed points. Dashed lines indicate unstable fixed points. H is the Hopf point.

We began by configuring the parameters so that both cell populations were nominally at rest in the absence of stimulation (*J* =0). This was done by choosing the connection weights (*w_ee_* = 12, *w_ei_* = 10, *w_ie_* = 10, *w_ii_* = 1) and firing thresholds (*b_e_*=1.75 and *b_i_*=2.6) so that the nullclines crossed near the left knee of the cubic nullcline (Fig 2D). The stable resting point for this configuration was *U_e_* = 0.12 and *U_i_* = 0.17. We then applied a constant stimulus (*J* =1) which induced a stable oscillation in *U_e_* and *U_i_* (Fig 2C). The limit cycle (black) is shown in Fig 2D. The time scales of excitation and inhibition (*τ_e_*=10, *τ_i_*=5) were adjusted so that the frequency of the oscillation was approximately 20 Hz, that being an appropriate time scale for neurons in visual cortex. Numerical continuation revealed that the limit cycle emerges via a supercritical Hopf bifurcation when the injection current exceeds the critical value *J* = 0.41 (Fig 2E). In this case, the limit cycle grows relatively smoothly with stimulus strength which we reasoned was an appropriate characteristic for obtaining a graded neural response to visual motion.

We then investigated the effects of inhibitory-surround coupling on the formation of endogenous waves in the spatially extended model (Fig 3A). In this case, the excitatory and inhibitory lateral projections both had Gaussian spatial profiles, *K_e_*(*x*) and *K_i_*(*x*), where the spread of the inhibitory coupling (*σ_i_*=0.15 mm) was three times broader than that of the excitatory coupling (*σ_e_*=0.05 mm). The spatial footprints of these projections spanned approximately 0.6 mm which is consistent with the anatomical span of pyramidal dendrites [53]. When combined, these excitatory and inhibitory profiles produced the classic ‘Mexican hat’ profile shown in Fig 3B (black). As anticipated, this configuration of lateral coupling elicited self-organized standing waves (Fig 3D) under spatially-uniform constant stimulation. Furthermore, the spatial frequency (2.5 cycles/mm) of the standing wave was predicted by the dominant spatial frequency of the Mexican hat, as we had previously seen in phase-based neural fields [52]. Nonetheless, slight variations in the selected pattern can occur on a trial to trial basis [54]. More importantly, we were able to transform the standing wave into a traveling wave (Fig 3E) by applying a small spatial shift (*δ*=0.02) to the excitatory coupling profile. The temporal frequencies of those traveling waves were typically 15 Hz, where negative frequencies indicate leftwards motion. Even though the shift in *K_e_*(*x δ*) was barely noticeable, it still produced a marked asymmetry in the Mexican hat (Fig 3C). Asymmetric coupling topologies are known to induce traveling waves in neural fields [25, 55]. In this case, the asymmetric Mexican hat operates like a spatial filter that responds maximally to waves that are phase-shifted to the right, so the wave travels to the left.

**Fig 3.**
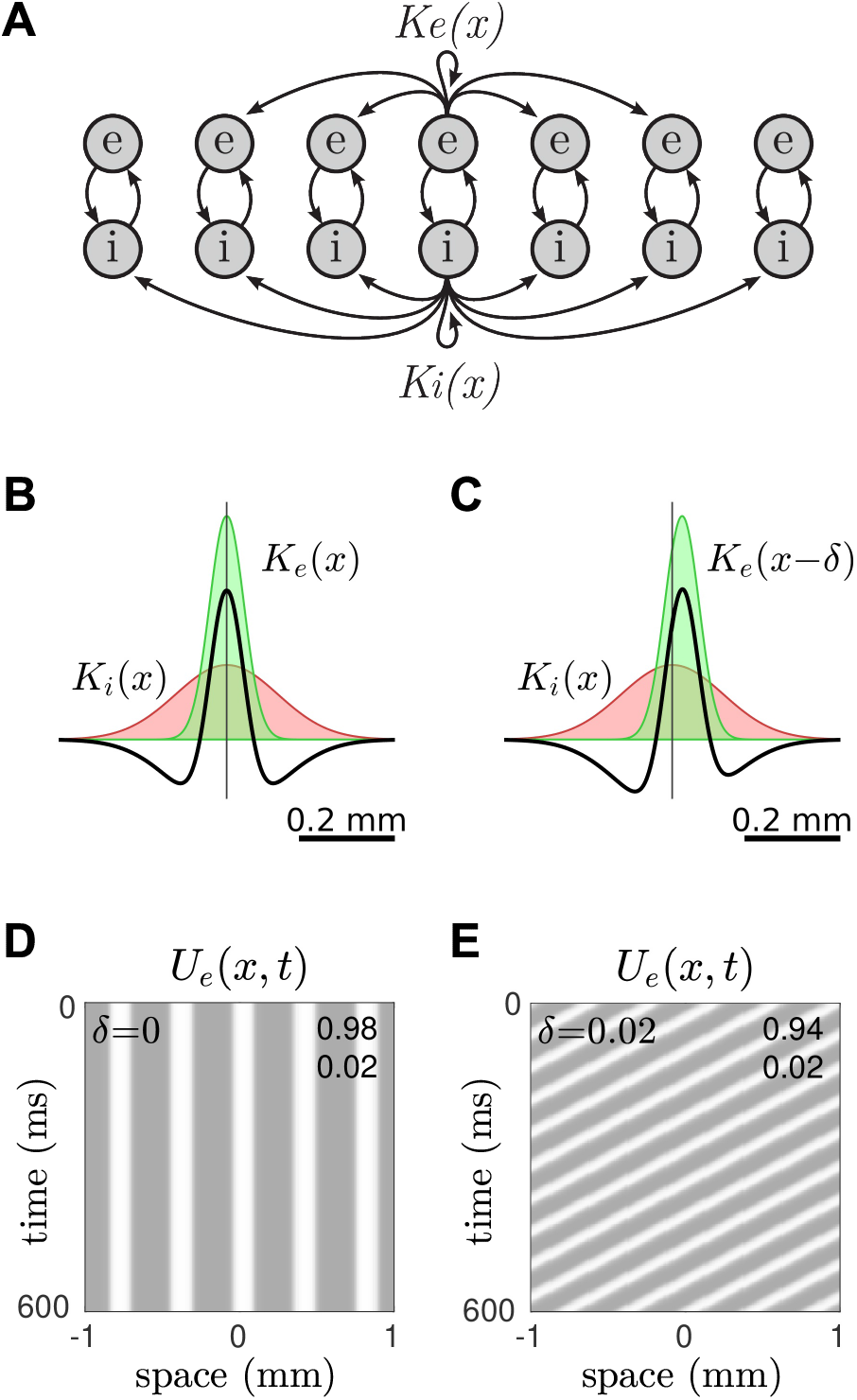
Endogenous traveling waves in the spatial model. A: Schematic of the lateral coupling. The spatial profiles of the excitatory and inhibitory projections are defined by *K_e_*(*x*) and *K_i_*(*x*) respectively. The inhibitory projections have the furthest reach. Only a few connections are shown. B: Symmetric Mexican hat coupling profile (black) constructed from symmetric Gaussian profiles for the excitatory cells (green) and inhibitory cells (red) respectively. C: Asymmetric Mexican hat obtained by shifting the excitatory coupling profile to the right by *δ*=0.02 mm. D: Stationary waves in the spatial model with symmetric lateral coupling, as per panel B. The gray scale indicates the mean firing rate of the excitatory cells. The minimum and maximum values are listed in the upper-right corner. E: Traveling waves in the spatial model with asymmetric lateral coupling, as per panel C.

Since the endogenous waves only emerged when the medium was stimulated, we hypothesized that it would respond preferentially to stimuli whose spatiotemporal signature best matched that of the endogenous wave. We tested this idea by stimulating the asymmetrically coupled medium with sinusoidal gratings that had identical spatial frequencies (*f_x_*=2.5 cycles/mm) but either moved in opposite directions (*f_t_*=−15 Hz versus *f_t_*=+15 Hz) or remained stationary (*f_t_*=0 Hz). As anticipated, the medium responded robustly to the stimulus whose frequency characteristics matched that of the endogenous wave (Fig 4A). However it also responded intermittently to the grating that moved in the opposite direction (Fig 4B) and the stationary grating (Fig 4C). For the case of motion in the opposite motion, the responses pulsated in time with the stimulus and appeared to lurch in the same direction but with occasional slips. The peak amplitude of the intermittent responses (*U_e,max_*=0.94 for the opposite grating; *U_e,max_*=0.95 for the stationary grating) actually exceeded that for the preferred stimulus (*U_e,max_*=0.89). The intermittent pulses evoked by the stationary grating were more regular. Nonetheless, as a putative motion detector, the proposed model (Fig 3A) failed to discriminate the preferred motion from the non-preferred motion.

**Fig 4.**
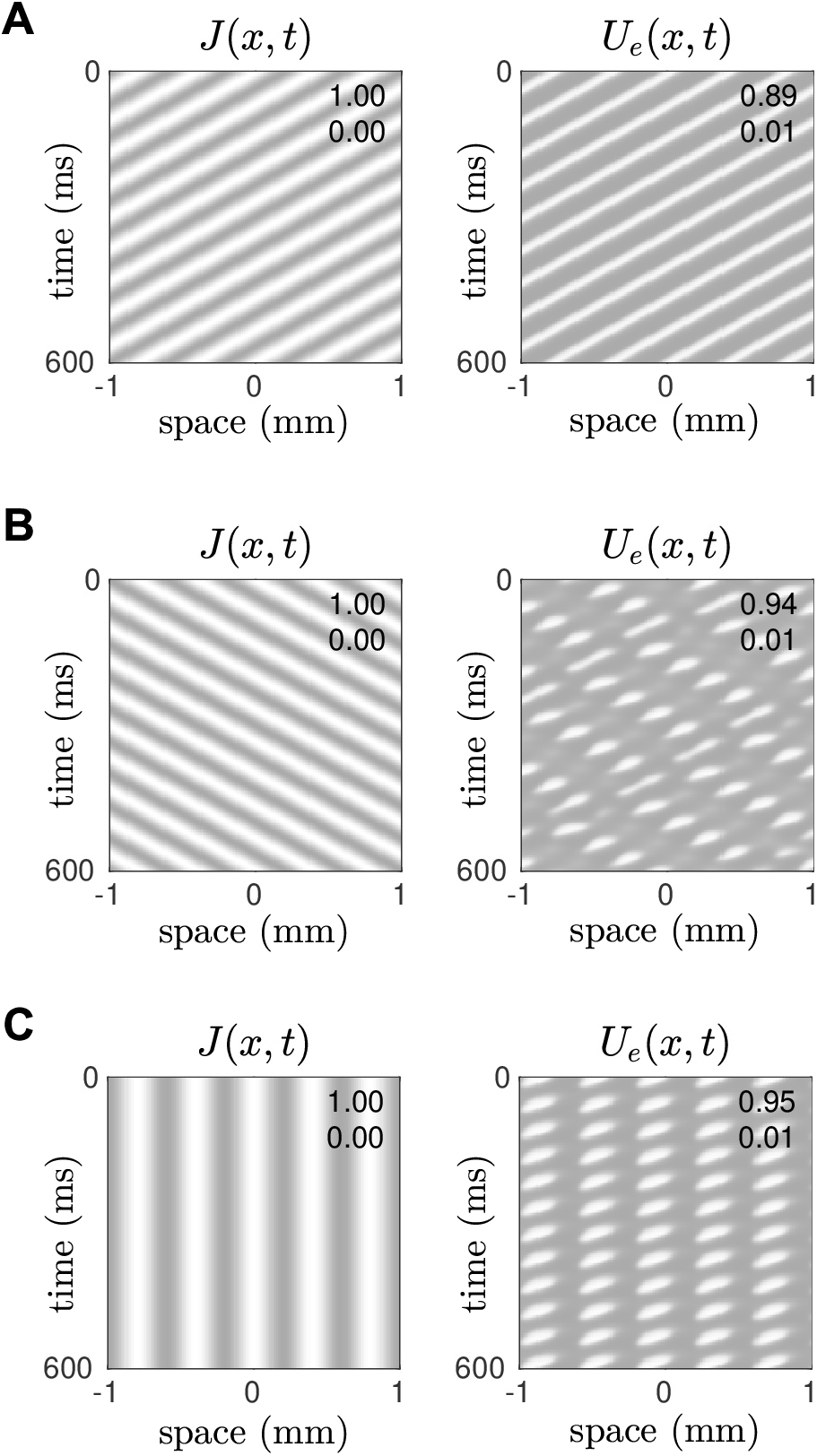
Effect of a moving grating stimulus on endogenous traveling waves. Here *J* (*x, t*) is the stimulus and *U_e_*(*x, t*) is the response of the medium. In all cases the medium is tuned (*δ*=0.02) for leftward propagating waves with a spatial frequency of *f_x_*=2.5 cycles/mm and a temporal frequency of *f_t_*=15 Hz. A: Case of a leftwards-moving grating whose spatiotemporal signature matches that of the endogenous waves. B: Case of a rightwards-moving grating (*f_t_*=−15 Hz). C: Case of a stationary grating (*f_t_*=0 Hz).

### The E-I-E model

We conjectured that this failure may be due to the model having insufficient degrees of freedom to accommodate the non-preferred motion signals. We therefore constructed a new model with an additional excitatory population that we call the E-I-E model (Fig 5A). The equations of this model were defined as,

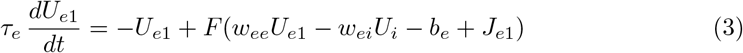

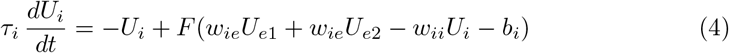

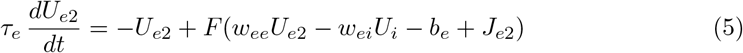

where *U*_*e*1_(*t*) and *U*_*e*2_(*t*) are the normalized firing rates of the two excitatory populations and *J*_*e*1_(*t*) and *J*_*e*2_(*t*) are their respective stimuli. All other parameters are the same as for Eq (1–2). The two excitatory populations in this model represent distinct assemblies of neurons that have the same firing characteristics but are not directly connected to one another. They can only interact via the common population of inhibitory neurons. The excitatory populations receive independent stimulation on the assumption that they are innervated by distinct incoming projections.

**Fig 5.**
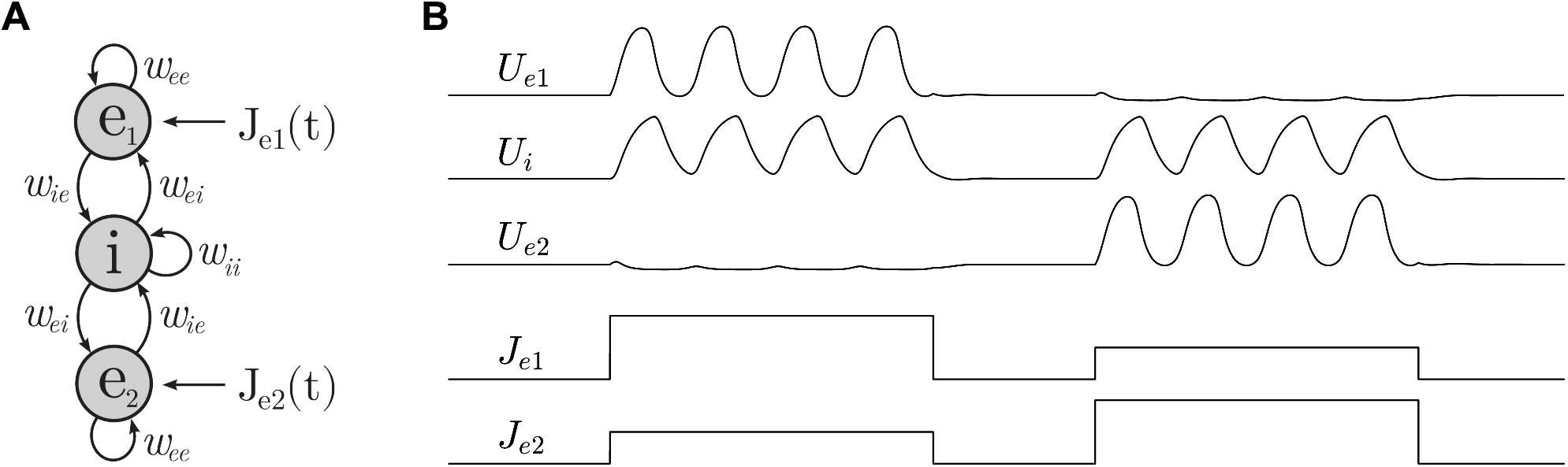
The E-I-E model. A: Schematic of the model. The excitatory populations *e*_1_ and *e*_2_ are not directly connected. B: Effect of differential stimulation of *e*_1_ and *e*_2_ where *J*_*e*1_ > *J*_*e*2_ in the first pulse and *J*_*e*1_ < *J*_*e*2_ in the second pulse. The responses in *U*_*e*1_ and *U*_*e*2_ are mutually exclusive and selective to the cell with the strongest stimulus.

The E-I-E model proved to be remarkably selective to differential stimulation. When stimuli of different magnitude (*J*_*e*1_≠*J*_*e*2_) were simultaneously applied to both populations, the responses in *U*_*e*1_ and *U*_*e*2_ always favored the population with the greatest input. Furthermore, those responses were mutually exclusive so that the ‘losing’ population was largely quiescent irrespective of how much it was stimulated (Fig 5B). This suggested that the E-I-E model robustly discriminates between incoming stimuli, even in the face of considerable ambiguity. We therefore analyzed the model’s behavior over a range of differential stimuli *J*_*e*1_ = *J* + Δ and *J*_*e*2_ = *J −* Δ which always favored population *e*_1_ (Fig 6C). In the analysis that follows, we used numerical continuation to follow the steady-state responses in *U*_*e*1_ and *U*_*e*2_ while ramping *J* and holding Δ fixed. We began with the case of ambiguous signals (Δ=0).

**Fig 6.**
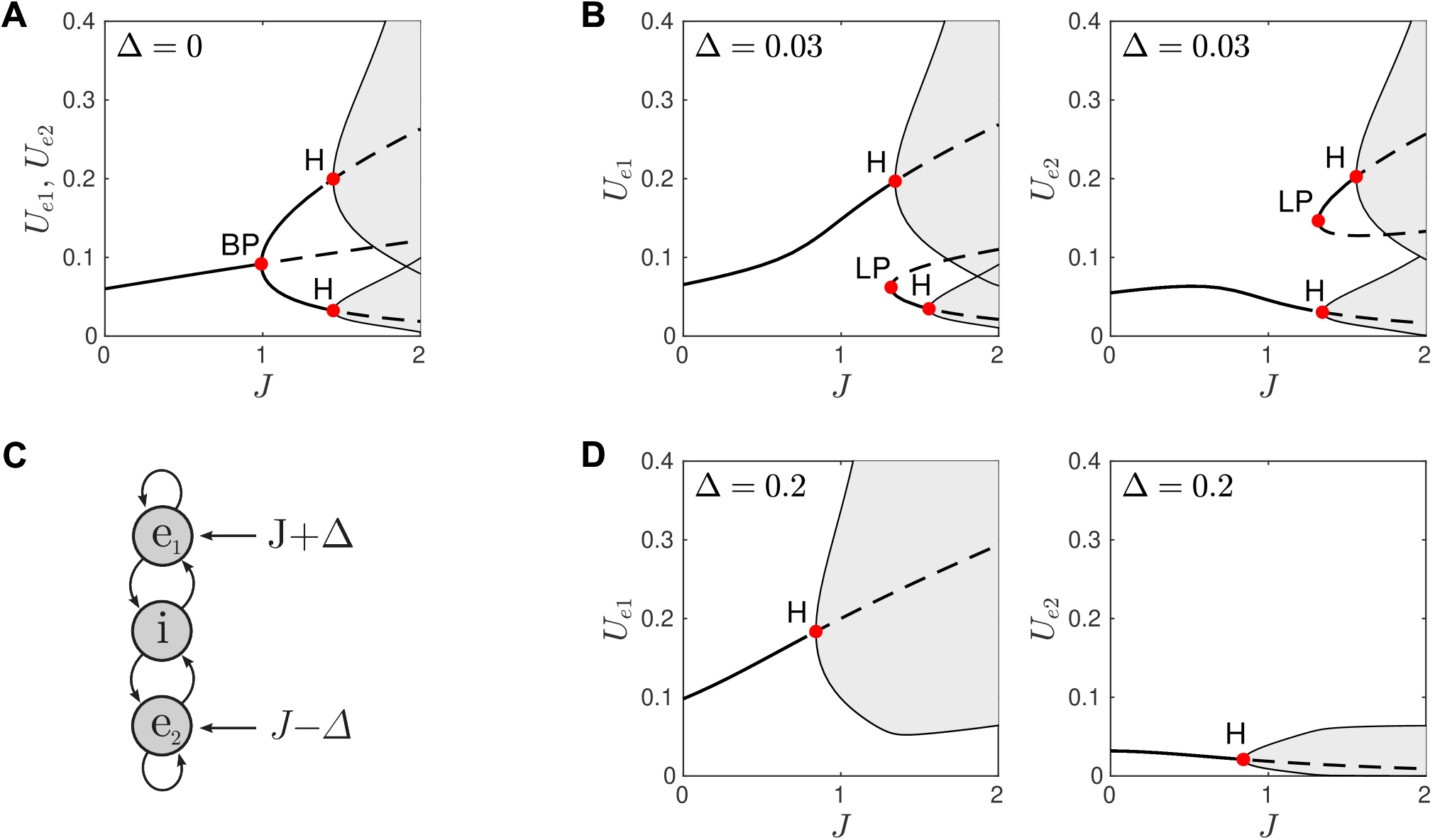
Bifurcations in the E-I-E model under differential stimulation. A: Responses to identical stimulation *J*_*e*1_ = *J* + Δ and *J*_*e*2_ = *J* − Δ where Δ=0. B: Responses to weakly biased stimulation (Δ=0.03). C: Schematic of the model. D: Responses to moderately biased stimulation (Δ=0.2). Solid lines indicate stable fixed points. Dashed lines indicate unstable fixed points. Shaded regions are the envelopes of limit cycles. BP is branch point. H is Hopf bifurcation. LP is limit point.

### Selective responses to ambiguous stimuli

Fig 6A shows the bifurcation diagrams for both *U*_*e*1_ and *U*_*e*2_ for the case of Δ = 0 where the diagrams are identical because of symmetry. For *J* < 1 the responses in *U*_*e*1_ and *U*_*e*2_ are monostable fixed points which are necessarily identical. Those fixed points diverge at *J* = 1 via a pitchfork bifurcation at the branch point (labeled BP). For *J* > 1 the steady states of *U*_*e*1_ and *U*_*e*2_ may follow either of the upper or lower branches of stable fixed points. Those selections are essentially at random but they are also mutually exclusive. Thus if *U*_*e*1_ selects the upper branch then *U*_*e*2_ selects the lower branch, and vice versa. The branch of identical fixed points (*U*_*e*1_=*U*_*e*2_) continues to exist for *J* > 1 but is unstable (dashed line) and forms a separatrix between the two branches of stable fixed points.

For *J* > 1.4 the fixed points lose stability via supercritical Hopf bifurcations (labeled H) that give rise to co-existing stable limit cycles (shaded). The ambiguous stimulus allows *U*_*e*1_ and *U*_*e*2_ to select either of those limit cycles. As before, those selections are mutually exclusive. So if *U*_*e*1_ selects the oscillation on the upper branch then *U*_*e*2_ selects the oscillation on the lower branch, and vice versa. The oscillations on the upper branch are much larger than those on the lower branch. We regard the winner of the competition between *e*_1_ and *e*_2_ to be the one that selects the branch of large oscillations.

So for ambiguous stimuli with *J* > 1.4, the winner is essentially selected at random but at least that selection is decisive. The separatrix between the upper and lower branches of solutions is the key to that selectivity because it forces the responses of *e*_1_ and *e*_2_ to self-segregate even though the stimuli (*J*_*e*1_=*J*_*e*2_) are identical.

### Selective responses to weakly biased stimuli

The selectivity of the E-I-E model is no longer random once the stimulus is biased (Δ = 0). Fig 6B shows the bifurcation diagrams for *U*_*e*1_ (left panel) and *U*_*e*2_ (right panel) for the case of weakly biased stimulation (Δ=0.03). The pitchfork bifurcation is replaced by an ‘imperfect’ bifurcation that has no branch point. For *J* < 1.3 the stable fixed points in *U*_*e*1_ and *U*_*e*2_ are both monostable. More importantly *U*_*e*1_ steadily increases with *J* whereas *U*_*e*2_ steadily decreases. This divergence in responses guarantees that *e*_1_ wins the competition — provided that the stimulus is ramped slowly from zero. The perceptual decision is robust for *J* > 1.34 where large oscillations emerge on the upper branch (left panel; upper H) and small oscillations emerge on the lower branch (right panel; lower H). The small oscillations in *U*_*e*2_ are negligible compared to the large oscillations in *U*_*e*1_.

However that outcome is not guaranteed when the stimulus is suddenly onset rather than slowly ramped. In that case, it is possible for *U*_*e*1_ and *U*_*e*2_ to select other stable states that co-exist for *J* > 1.32 where a pair of stable and unstable fixed points emerge from the limit point (LP). This minor branch of stable fixed points itself gives way to stable oscillations for *J* > 1.56. For *U*_*e*1_ those oscillations are small (left panel; lower H) and for *U*_*e*2_ those oscillations are large (right panel; upper H). If *e*_2_ happens to select that large-amplitude oscillation then it wins the competition and the perceptual decision is a false positive. The mistake is forgivable given that Δ = 0.03 is a very weak bias in this case.

The potential for false positives is due to the existence of the limit point (LP). It is a remnant of the branch point (BP in Fig 6A) that is lost when Δ ≠ 0 transforms the pitchfork bifurcation into an imperfect bifurcation. The position of the limit point is governed by the size of the bias in the stimulus. Increasing Δ > 0 shifts the limit point towards higher *J*. If the bias is large enough then it effectively eliminates the false positives by shifting the limit point beyond the operating range of *J*.

### Selective responses to strongly biased stimuli

Fig 6D shows the bifurcation diagrams for *U*_*e*1_ (left panel) and *U*_*e*2_ (right panel) for the case of strongly biased stimuli (Δ=0.2). The limit point has been shifted beyond *J* > 2 and the remaining steady states are all monostable. Hence *e*_1_ is guaranteed to win the competition for *J* > 0.84 where a large oscillation emerges in *U*_*e*1_ and a small corresponding oscillation emerges in *U*_*e*2_. Once again, the small oscillations in *U*_*e*2_ are negligible compared to the large oscillations in *U*_*e*1_. The strong bias (≈20% of baseline) thus ensures that the correct response is always selected.

### The spatial E-I-E model

Returning to the problem of motion discrimination, we constructed a spatial variant of the E-I-E model where the profiles of the lateral projections in the excitatory layers were shifted in opposite directions (*δ* = ±0.02 mm) while the profile of the inhibitory projections remained symmetric (Fig 7A). This coupling topology produced asymmetric Mexican hat profiles for both the upper and lower layers of the model (Figs 7B,C). The spatiotemporal stimulus *J* (*x, t*) was applied identically to both of the excitatory layers in this model. We hypothesized that the opposing phase shifts in the lateral coupling profiles would impel waves in the top layer to travel leftwards and those in the bottom layer to travel rightwards. While the external stimulation would serve as a bias that favored the layer which best matched the spatiotemporal signature of the stimulus. We reasoned that the selective response properties observed in the E-I-E point model would also apply to spatiotemporal activity patterns in the spatial model.

**Fig 7.**
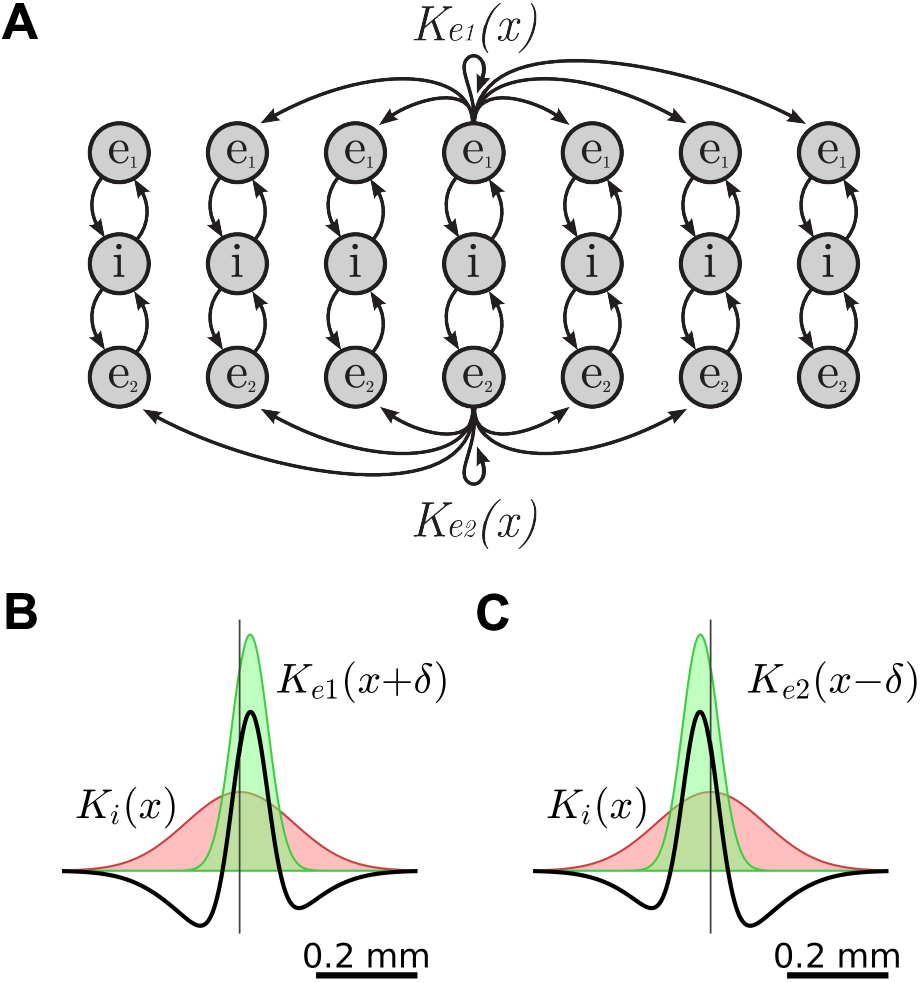
The spatial E-I-E model with asymmetric lateral coupling. A: Schematic of the model where the spatial profiles of the excitatory projections, *K_e_*_1_(*x*) and *K_e_*_2_(*x*), are shifted in opposite directions. The lateral inhibitory projections (not shown) remain symmetric. B: Asymmetric Mexican hat constructed from *K_e_*_1_(*x*+*δ*) and *K_i_*(*x*) where *δ*=0.02 mm. C: Asymmetric Mexican hat constructed from *K_e_*_2_(*x−δ*) and *K_i_*(*x*). Note the opposing phase shifts in the Mexican hat profiles.

We tested this concept by simulating the spatial E-I-E model with the same drifting gratings that we used in Fig 4 and found that the excitatory layers of the model were exquisitely selective to the direction of the moving stimulus. Moreover we saw no false responses to motion in the opposite direction. Fig 8A shows the response to a leftward moving grating whose spatial (*f_x_*=2.5 cycles/mm) and temporal (*f_t_*= − 15 Hz) frequencies match those of the endogenous wave in the top layer of excitatory neurons, represented by *U*_*e*1_(*x, t*). The responses in *U*_*e*1_(*x, t*) spanned the majority of the variable’s dynamic range (0.01<*U*_*e*1_<0.89) whereas that in *U*_*e*2_(*x, t*) was very much suppressed (0 < *U*_*e*2_ < 0.01). We interpreted the overwhelmingly dominant activity of the *e*_1_ layer as a robust perceptual response to leftwards motion.

**Fig 8.**
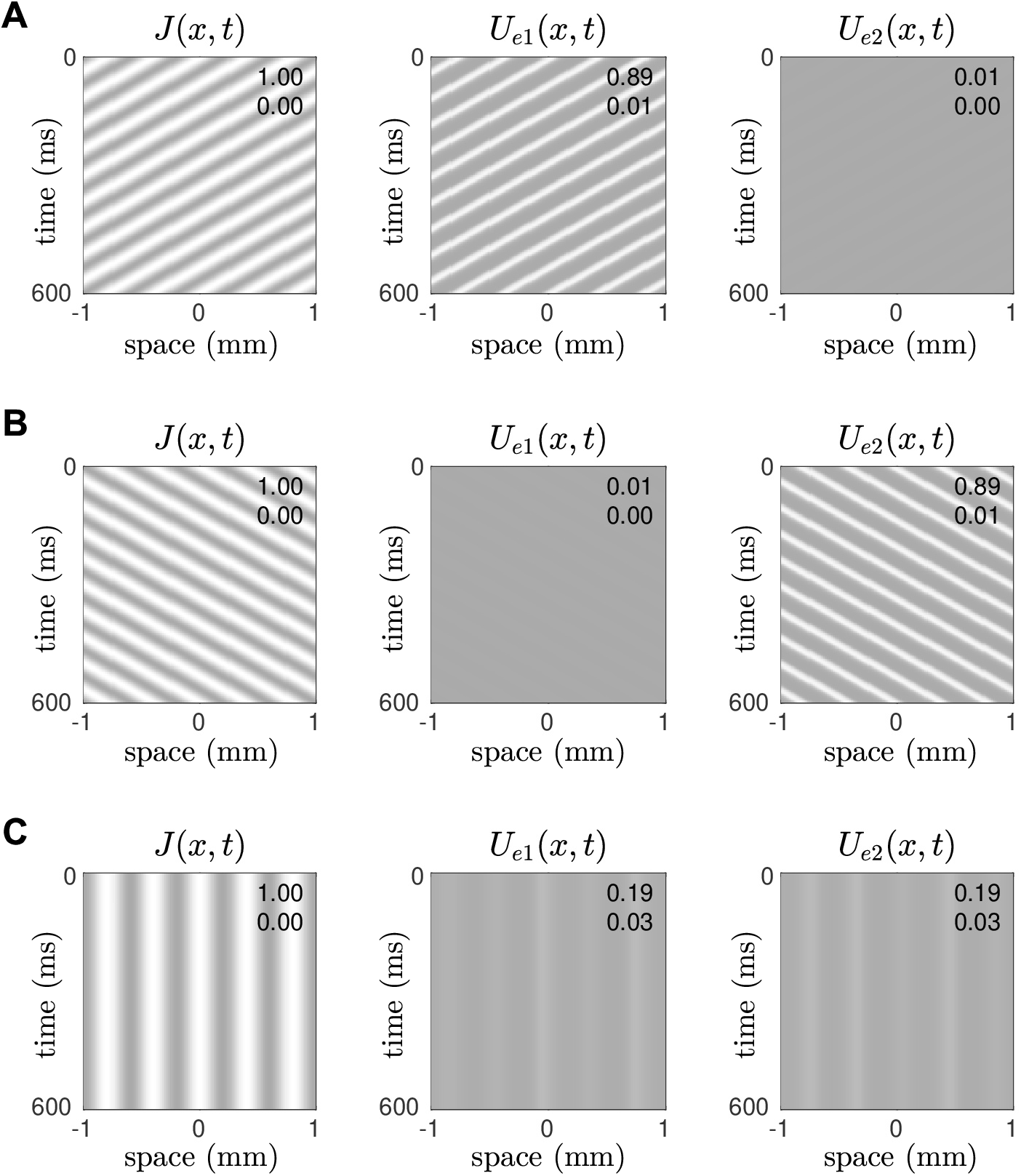
Direction-selective responses in the spatial E-I-E model. The cells in layer *e*_1_ were tuned to leftward motion (*δ*=+0.02) and those in layer *e*_2_ were tuned to rightward motion (*δ*=−0.02). The external stimulus *J* (*x, t*) was applied identically to both layers. Their spatiotemporal responses are *U*_*e*1_(*x, t*) and *U*_*e*2_(*x, t*). A: Case of a leftwards moving grating (*f_x_*=2.5 cycles/mm, *f_t_*=15 Hz) which resonates with the endogenous wave in *U*_*e*1_. B: Case of a rightwards moving grating (*f_x_*=2.5 cycles/mm, *f_t_*=−15 Hz) which resonates the endogenous wave in *U*_*e*2_. C: Case of a stationary grating (*f_x_*=2.5 cycles/mm, *f_t_*=0 Hz) which does not resonate with either.

The symmetric result was also observed for a rightward moving grating whose spatial (*f_x_*=2.5 cycles/mm) and temporal (*f_t_*=−15 Hz) frequencies match those of the endogenous wave in the bottom layer of excitatory neurons, represented by *U*_*e*2_(*x, t*). In that case, the *e*_2_ layer gave the dominant response and the *e*_1_ layer was suppressed (Fig 8B). The model responded as equally robustly to rightwards motion as it did to leftwards motion in these two test cases. The response to stationary gratings (Fig 8C) was also pleasing as both layers *e*_1_ and *e*_2_ exhibited suppressed responses (0.03*<U*<0.19) with no temporal oscillations. Such an outcome is the spatiotemporal analogy of the diverging branches of fixed point solutions in the point model under ambiguous stimulation (Fig 6A). Here that divergence is expressed as subtle differences in the spatial patterns in *U*_*e*1_(*x, t*) and *U*_*e*2_(*x, t*) where the presence of a stationary pulse in one pattern tends to suppress a corresponding pulse in the other. This is evident in the substantial range of the point-wise differences between the two patterns, −0.14 < *U*_*e*1_(*x, t*) *− U_e_*_2_(*x, t*) < 0.14.

### Tuning curves

The previous simulations (Fig 8) demonstrated robust discrimination between leftward and rightward motion in specific test cases. We sought to generalize those findings by quantifying the responses of *U*_*e*1_(*x, t*) and *U*_*e*2_(*x, t*) to stimulus gratings with a range of spatial (0 < *f_x_* < 15) and temporal (−15 < *f_t_* < 15) frequencies.

The temporal frequency tuning curve (Fig 9A) was obtained by varying *f_t_* while holding the spatial frequency of the stimulus grating fixed at *f_x_* = 2.5 cycles/mm. It plots the maximal responses in *U*_*e*1_(*x, t*) and *U*_*e*2_(*x, t*) over the long term. The individual tuning curves for *U*_*e*1_ (dotted line) and *U*_*e*2_ (solid line) exhibit dramatic separation whereby *e*_1_ responds predominantly to leftward moving gratings (*f_t_* < 0) and *e*_2_ responds predominantly to rightward moving gratings (*f_t_* > 0). Furthermore the responses are sharply constrained to the 5–28 Hz frequency band which is why the stationary grating (*f_t_* = 0) did not elicit a strong response in either *U*_*e*1_ or *U*_*e*2_ (Fig 8C).

**Fig 9.**
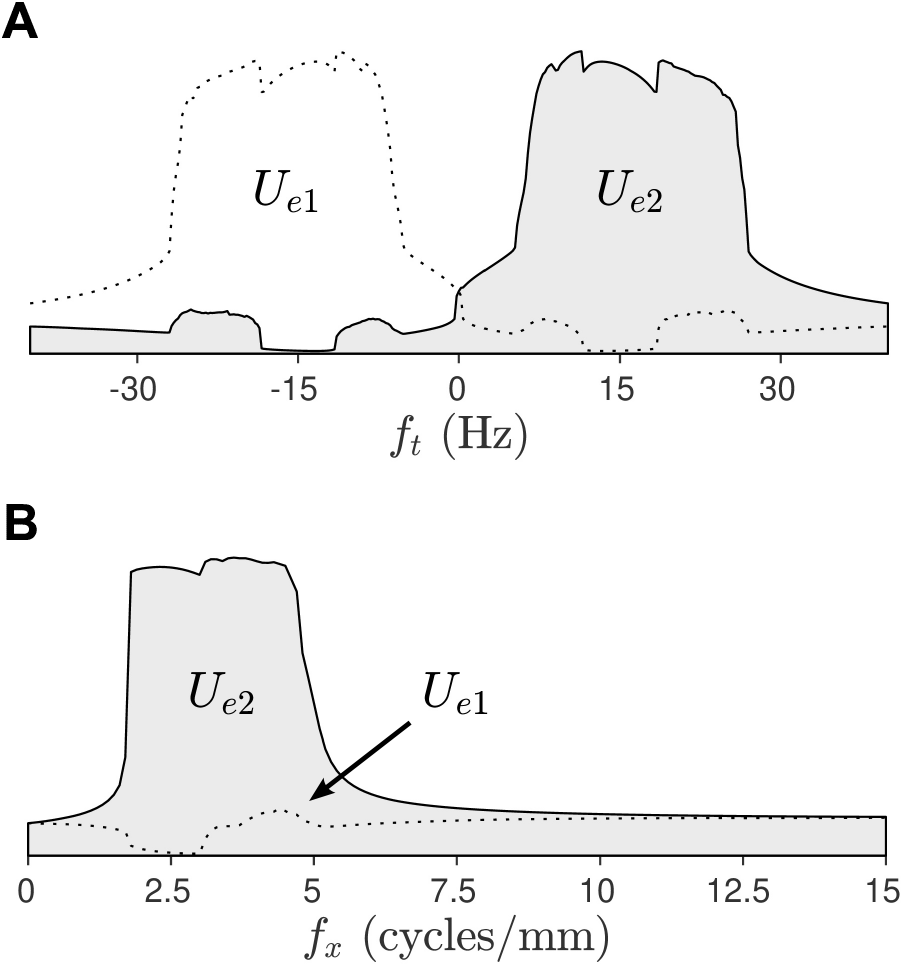
Tuning curves for the spatial E-I-E model. A: Temporal frequency tuning curve showing the maximal responses in *U*_*e*1_ and *U*_*e*2_ for stimulus gratings with temporal frequencies −40 < *f_t_* < 40 Hz where negative frequencies correspond to leftward motion. The spatial frequency of the grating is fixed at *f_x_* = 2.5 cycles/mm. B: Spatial frequency tuning curve showing the maximal responses to gratings with spatial frequencies 0 < *f_x_* < 15 cycles/mm. In this case the temporal frequency is fixed at *f_t_* = 15 Hz.

The spatial frequency tuning curve (Fig 9B) was similarly obtained by varying *f_x_* while holding the temporal frequency fixed at *f_t_* = 15 Hz which corresponds to rightwards motion. For this particular temporal frequency, the tuning curve for *U*_*e*2_ (solid line) is strongly selective to gratings with spatial frequencies 1.7 < *f_x_* < 5.0 cycles/mm. The response band is also remarkably sharp. Whereas the response for *U*_*e*1_ (dotted line) is attenuated at all spatial frequencies because it is tuned to motion in the opposite direction. The converse behavior is observed for leftward moving gratings (*f_t_*=−15 Hz).

## Discussion

Our model demonstrates how neurons in the visual cortex can exploit endogenous background wave activity to compute non-separable spatiotemporal receptive fields without resort to transmission delays. The proposed mechanism relies on the predisposition of the cortical tissue to generate traveling waves of activity whose speed and direction are determined by the lateral coupling topology. The waves act as spatiotemporal filters that selectively amplify those stimuli that have similar space-time signatures to the wave — after retinotopic mapping of the visual field onto the cortex. The selectivity is further enhanced by competition between waves that travel in opposite directions. That competition is mediated by the common population of inhibitory cells which provide negative feedback to the excitatory cells of the opposing waves. The dynamics of the E-I-E assembly are such that compromise solutions between competing stimuli are inherently unstable, leading to winner-take-all perceptual decisions. In the case of the point model, the competition is won by the excitatory cell with the stronger point stimulus. In the case of the spatial model, it is won by the excitatory cells whose endogenous wave pattern resonates most with the spatiotemporal stimulus. The competition suppresses responses that may arise in the opposing detector due to partial resonance of the stimulus with the opposing wave. Opponency is thus an effective neural strategy for suppressing false-positives in otherwise imperfect detectors.

Our proposal offers new theoretical insights into how direction-selectivity can arise in the visual cortex through the collective behavior of neurons therein. The endogenous wave activity imposes a spatiotemporal arrangement on the background neural activity that predisposes it to the preferred stimulus. Direction-selectivity is therefore not a property of a single neuron but is inherited from the collective behavior of many neurons. If such a neuron were to be isolated from its neighbors then it would immediately lose its spatiotemporal response properties.

Our model also suggests that excitation and inhibition should be expected to co-vary greatly in response to the preferred stimulus. This is consistent with invasive recordings of the dendritic currents in the primary visual cortex of anesthetized cats by Priebe and Ferster [56]. They found that the excitatory and inhibitory currents co-vary at different phases and that the peaks of the inhibitory currents were maximal for the preferred stimulus rather than the null stimulus — in contradiction to the standard model [56]. In our model, inhibition rises and falls because it is a mechanism of oscillation rather than a mechanism of stimulus suppression. The stimulus evokes maximal oscillations in the preferred excitatory cells while suppressing activity in the opposing excitatory cells. The inhibitory cells respond maximally either way except when both populations of excitatory cells are silent.

The importance of opponency has been demonstrated in many perceptual studies [57]. In theoretical models it is typically portrayed as a hypothetical subtraction between the outputs of neurons with opposing preferences. In the case of the E-I-E circuitry, the mechanism of opponency is inherent within the recurrent inhibitory feedback. It plays a dual role in driving the oscillatory dynamics as well as selecting the winning response. Without opponency, the simpler E-I model fails to discriminate against the non-preferred stimuli because it lacks the degrees of freedom to accommodate other scenarios. The opposing E-I-E circuitry, on the other hand, has enough degrees of freedom to allow the winner to alleviate the frustration of the loser.

Zhang [58] previously proposed a similar double-ring network with asymmetric lateral coupling for head-direction tuning cells in the hippocampus. In that model, the position of the head was encoded by a bump attractor that was continuously shifted one way or the other to integrate signals from proprioceptors in the head. The motion of the bump was driven by asymmetric coupling that was modulated in time by the proprioceptors for clockwise and anti-clockwise movement. The double rings operated in opposition in the sense that they pushed and pulled the bump in opposite directions. However the goal of that mechanism was to integrate movements from opposing proprioceptors rather than to suppress competing perceptual decisions, as is the goal of our model. Curtu and Ermentrout [59] showed that bump attractors can also be made to travel with slow negative feedback rather that asymmetric lateral coupling. In the absence of a stimulus, the direction of travel is determined by initial conditions. It is likely that a moving stimulus could force the bump to travel in the same direction but it is not clear how that mechanism could be used to discriminate between motion in opposite directions.

The E-I-E model has some interesting properties that make it an effective neural circuit for resolving competing responses, potentially in any sensory domain. The symmetry of the circuit means that symmetric solutions to ambiguous stimuli do exist but those solutions lose stability when the baseline stimulus exceeds a critical threshold. The unstable symmetric solutions thus act as a separatrix between co-existing stable solutions which are dominated by each of the opposing excitatory populations. Whether those stable solutions are fixed points or limit cycles depends largely upon the choice of *τ_e_* and *τ_i_*. Oscillations were crucial to the spatiotemporal signature of visual motion in the present study. However that need not be the case for other sensory domains where fixed points may be more appropriate.

Like many computational theories of vision, our proposal relies on the retinotopic mapping between the visual field and the cortex to preserve the geometric relationship between the stimulus and the endogenous neural activity. For simplicity, we assumed a one-to-one mapping between visual and cortical coordinates whereas the anatomical mapping is actually a log-polar relationship [60]. We anticipate that the use of log-polar retinotopic mapping would likely extend our results to the motion of rotating spirals, radial spokes and expanding rings [42] although our model has yet to be tested in two spatial dimensions. To do so requires constructing opposing pairs of motion detectors to accommodate multiple directions of motion. One approach would be to construct an asymmetric coupling function that rotates continuously in space to accommodate the different directions. Bressloff [61] has previously used that approach to model the orientation preferences of anatomical hypercolumns in primary visual cortex in relation to geometric visual hallucinations. In the case of visual motion detection, we believe that having a small number opposing detectors aligned along the cardinal axes of motion would likely suffice. Further research is required to determine how such an arrangement would effect the tuning curves.

## Methods

The equations for the spatial Wilson-Cowan model (Fig 3A) were defined as,

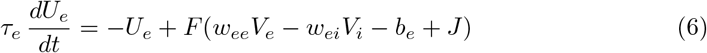

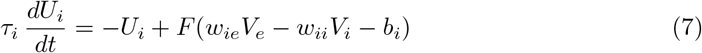

where *U_e_*(*x, t*) and *U_i_*(*x, t*) are the spatiotemporal firing rates of the excitatory and inhibitory neural populations with *x ∈* R^1^. The sigmoidal function,

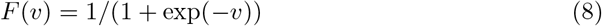

defined the firing rate of each cell population in response to the net input *v*. That input comprised of the spatially weighted activity *V_e_*(*x, t*) and *V_i_*(*x, t*) from nearby excitatory and inhibitory cells. The spatial summation,

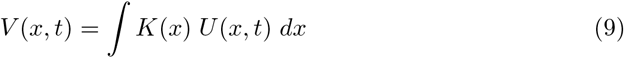

was computed by convolving the spatial activity in *U* (*x, t*) with the Gaussian kernel,

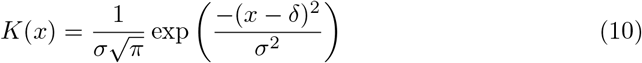

where *σ* is the spatial spread parameter and *δ* is a spatial shift parameter. The spatial shift was only applied to the excitatory cells. The connection weights *w* are scalar constants where *w_ei_* denotes the connection from an inhibitory population to an excitatory population. Parameters *b_e_* and *b_i_* represent the firing thresholds for the excitatory and inhibitory populations. Parameters *τ_e_* and *τ_i_* are the time constants of excitation and inhibition. All parameter values are listed in Table 1.

**Table 1.** Parameters of the model.

| Parameter | Description |
| --- | --- |
| $U_e(x, t)$ | Excitatory continuum firing rate |
| $U_i(x, t)$ | Inhibitory continuum firing rate |
| $J(x, t)$ | Spatiotemporal stimulus |
| $K(x)$ | Spatial coupling kernel |
| $\sigma_e = 0.05$ | Spread of excitation (mm) |
| $\sigma_i = 0.15$ | Spread of inhibition (mm) |
| $\delta = 0.02$ | Gaussian spatial shift (mm) |
| $w_{ee} = 12$ | Coupling weight ($e$ to $e$) |
| $w_{ei} = 10$ | Coupling weight ($i$ to $e$) |
| $w_{ie} = 10$ | Coupling weight ($e$ to $i$) |
| $w_{ii} = 1$ | Coupling weight ($i$ to $i$) |
| $b_e = 1.75$ | Excitatory threshold |
| $b_i = 2.6$ | Inhibitory threshold |
| $\tau_e = 5$ | Excitatory time constant (ms) |
| $\tau_i = 10$ | Inhibitory time constant (ms) |
| $\alpha$ | Stimulus amplitude |
| $f_x$ | Spatial frequency (cycles/mm) |
| $f_t$ | Temporal frequency (cycles/ms) |
| $dx = 0.01$ | Spatial step (mm) |

Similarly, the equations for the spatial E-I-E model (Fig 5A) were defined as,

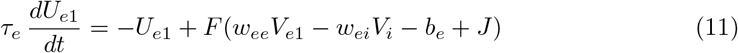

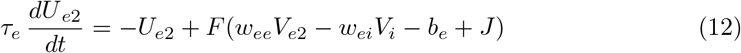

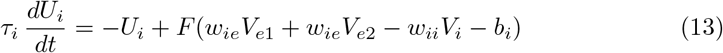

where *U*_*e*1_(*x, t*) and *U*_*e*2_(*x, t*) are the firing rates of the excitatory cells and *U_i_*(*x, t*) is the firing rate of the inhibitory cells.

Visual stimulation was represented by the spatiotemporal signal *J* (*x, t*) which was applied to the excitatory cells only. It was defined as a sinusoidal moving grating,

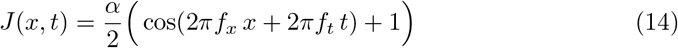

where *α* is the amplitude of the grating and *f_x_* and *f_t_* are its spatial and temporal frequencies.

### Numerical Simulation

The forward models were simulated using Version 2019a of the Brain Dynamics Toolbox [62, 63] running in Matlab R2019b. The differential equations were integrated forward in time using the `ode23` solver with variable time steps and error tolerances of `AbsTol=1e-6` and `RelTol=1e-6`. The numerical continuation was performed using Matcont [64, 65] version 7p1 with the default tolerances. The step size for branch of equilibrium points was limited to `MaxStepSize=0.1`. The step size for the branch of limit cycles was limited to `MaxStepSize=0.5`.

## Acknowledgments

This work is based on a conference paper [66] that was originally presented at the First International Workshop on Computational Models of the Visual Cortex (CMVC) at Columbia University and published in the Proceedings of the 9th EAI International Conference on Bio-inspired Information and Communication Technologies (BICT 15), New York City.

## Author Contributions

SH and BE contributed to the conception and design of the study; SH wrote the software and performed the simulations; SH wrote the first draft of the manuscript and BE revised it for intellectual content. Both authors contributed to subsequent revisions of the manuscript and they both read and approved the final version.

## Notes

#### Summary of Updates

Added a reference to Lien and Scanziani (2018).

